# Muscle Loss as a Foundational Step in the Development and Evolution of the Turtle Shell

**DOI:** 10.64898/2026.06.16.732726

**Authors:** William Foster, Paul Gensbigler, Max Blecher, Andre Forjaz, Biyi Li, Valentina Matos-Romero, Chulan Kwon, Ashley Kiemen, Tyler Lyson, Gabriel S. Bever

**Affiliations:** Center for Functional Anatomy and Evolution, Johns Hopkins University School of Medicine, 1830 East Monument Street, Baltimore, MD 21287, USA; Department of Pathology, Sol Goldman Pancreatic Cancer Research Center, Johns Hopkins University School of Medicine, 600 N. Wolfe Street, Baltimore, MD 21287, USA; Department of Chemical & Biomolecular Engineering, Johns Hopkins University, 3400 N. Charles Street, Baltimore, MD 21218, USA; Department of Biomedical Engineering, Johns Hopkins University School of Medicine, 720 Rutland Avenue, Baltimore, MD, 21205, USA; Department of Oncology, Johns Hopkins University School of Medicine, 600 N. Wolfe Street, Baltimore, MD 21287, USA; Department of Earth Sciences, Denver Museum of Nature & Science, 2001 Colorado Boulevard, Denver, CO 80205, USA

## Abstract

Modern biodiversity is built on a series of disparate body plans whose origins are obscured by deep time. Our most direct source of clarifying data is the fossilized skeleton, where morphology reflects an evolving functionality realized through the development of associated tissues. We apply this dualistic perspective to the shelled body plan of turtles whose Paleozoic initiation is marked by a derived relationship between ribs and dermis (Lyson & Bever, 2020). Current developmental models remain in conflict with an increasingly informative fossil record, suggesting critical steps remain unrecognized. Here we explore the hypothesis that the breakdown of rib-spanning muscles—an evolutionary transformation mirrored in embryogenesis—is one such step. Multi-modal imaging of turtle embryos, including a novel application of histology-based deep learning (Kiemen et al., 2022; Matos-Romero et al., 2025; Forjaz et al., 2026), establishes intercostal muscle degradation as preceding turtle-specific rib development and highlights the heuristic power of 3D, whole-embryo analysis (Forjaz et al., 2026). Quantified divergence from mouse pinpoints the timing and tempo of this organized, apoptotic breakdown. Initial evidence suggests an associated non-pathological inflammatory response, which has been shown capable of driving evolutionarily stable hyperossification (Rashid et al., 2023). These patterns support trunk muscles as a critical signalling centre whose ontogenetic loss set the phylogenetic stage for a morphogenetic transformation remarkable in a non-metamorphic species.

## Main Text

The iconic shelled body plan of turtles, with dorsal carapace and ventral plastron, represents a radical rearrangement and hyperossification of the body wall that serves as a natural experiment in the morphogenetic potential of highly conserved vertebrate tissues (Burke, 1989; Gilbert et al., 2001; Kuratani et al., 2011; Lyson et al., 2013; Moustakas-Verho et al., 2017). Resolving what steps constitute this ‘turtle experiment’ is a long-standing problem in comparative biology and one that still ranks among its greatest intellectual challenges (MacCord et al., 2015; Wagner, 2015). The last quarter century has witnessed remarkable progress on the developmental underpinnings of turtle anatomy, as well as its deep evolutionary history as preserved in an increasingly informative fossil record (Schoch & Sues, 2020; Lyson & Bever, 2020). Both datasets grant foundational importance to a thoracic series of uniquely widened, T-shaped ribs and to ‘axial arrest’. The latter being the process through which the developing ribs grow laterally to give turtles their disc-like form and to penetrate a turtle-unique signalling centre along the dorsal flank called the carapacial ridge (CR) (Burke, 1989).

The role of the CR in patterning axial arrest is controversial but likely involves an FGF feedback-loop with the distal rib (Loredo et al., 2001; Cebra-Thomas et al., 2005; but see Nagashima et al., 2007; Kuratani et al., 2011; Hirasawa et al. 2013). The result is a diversion of the turtle rib from its ancestral, ventral trajectory into the lateral plate mesoderm and its retention as a fully primaxial structure (i.e.; it remains on the dorsal, primaxial side of the lateral somitic frontier [Burke and Nowicki, 2003; Nowicki et al., 2003]). Axial arrest and CR penetration by the rib is followed by BMP-mediated metaplastic ossification of rib-adjacent dermis, producing a compound structure (i.e., costal) with a T-shaped, cross-sectional form (Lyson et al., 2013; Rice et al., 2015) (Fig. 1B). This largely postnatal phenomenon continues until neighbouring ribs become suturally locked in what is now a maturing carapace. From this developmental progression flows the hypothesis that the CR directs axial arrest and costal formation and is a key innovation whose own origin extends to the earliest history of the turtle stem lineage. The problem is that while the phylogenetically earliest of the hypothesized stem turtles—middle Permian (c.260mya) *Eunotosaurus africanus* (Lyson et al., 2010, 2013, 2014, 2016; Bever et al., 2015) and late Triassic (c.240mya) *Pappochelys rosinae* (Schoch and Sues, 2015, 2018)—exhibit T-shaped ribs, these ribs clearly enter the ventral body wall and thus lack axial arrest. This contradiction suggests axial arrest and costal formation are patterned independently and that key developmental dynamics remain unrecognized or unappreciated. Here, we explore this possibility by tracking body wall embryogenesis in a collection of amniotes (turtle, chick, alligator, and mouse) using multiple three-dimensional (3D) imaging modalities including the deep-learning software CODA (Supplementary Figs. 1-3), which is designed to identify, quantify and analyse a multitude of cell and tissue types in serially sectioned histology(Kiemen et al., 2022; Gensbigler et al., 2025). From these data, we derive the hypothesis that degradation and loss of the intercostal musculature (ICM) is not a secondary product of the shell-forming process but a foundational event in the derived patterning of the body wall that helped move turtles along their unlikely evolutionary path.

**Fig 1.**
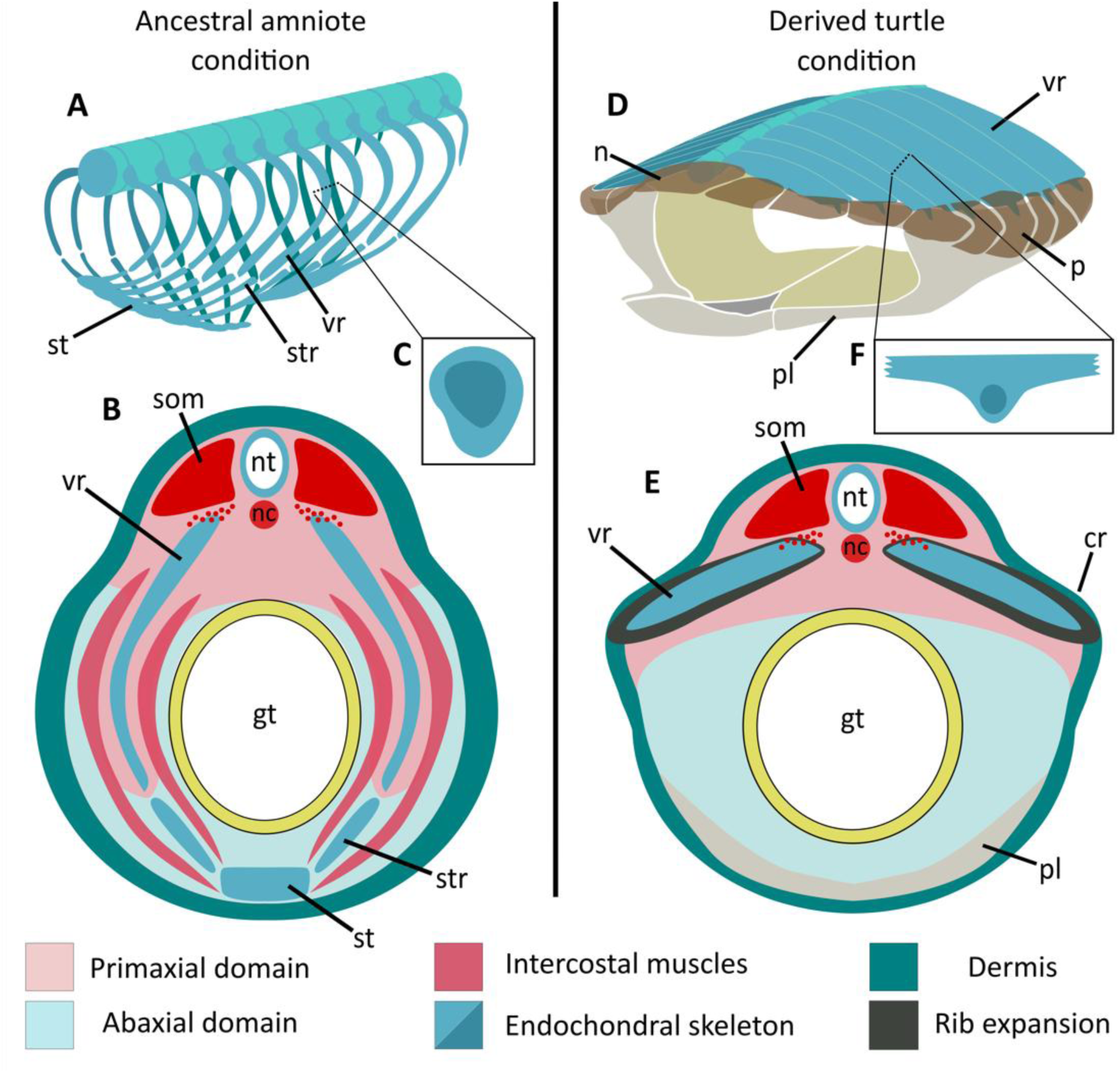
Derived nature of the secondary body wall in the modern turtle body plan. (A, B) Schematic representation of the ancestral amniote body wall in left anterolateral view and transverse section, respectively. Sub-ovoid (C) endochondral ribs grow into the ventral body wall, defining intercostal spaces that are filled with a highly organized, layered musculature. These features together produce a flexible but strong thoracic wall whose movement facilitates the volumetric and pressure dynamics driving lung ventilation. Turtle ribs (D-F) do not enter the ventral body wall but instead grow laterally into a turtle-unique signalling center called the carapacial ridge. Secondary ossification of the intercostal spaces gives the ribs a T-shaped cross-sectional form that eventually serves as the structural framework for the inflexible upper shell. **Abbreviations:** cr, carapacial ridge; gt, gut tube; nc, notochord; nt, neural tube; pl, plastron; som, somites; st, sternum; str, sternal rib; vr, vertebral rib.

### Early Morphogenesis of the Turtle Body Wall

The early features of secondary body wall formation, which appear just prior to Greenbaum stage (Greenbaum, 2002) (G) 12 turtle embryo (Fig. 2A), are highly similar among sampled amniotes and thus likely conserved across this >300 million-year (my) radiation (Fig. 2B-D). Somitic delamination in the thoracic series begins cranially as a series of wave-like fronts with sclerotomal and myotomal cell populations advancing into the primary body wall. By G12, myotomal delamination is evident along the entire length of the thorax, with anterior segments maintaining an advanced state of differentiation that includes the clear presence of skeletal muscle fibres. These fibres are densely packed and insert onto newly established and uncurved rib anlagen as ICM. Distal to the developing ribs, the G12 fibres are less dense and more loosely arranged. In this distal region, the successive myotomal fronts form a continuous plate that by G13 spans the craniocaudal length of the thorax but remains dorsal to somatopleure and coelom (Supplementary Fig. 4). The conserved nature of these early myotomal dynamics is further evidenced by the similar rates with which the trunk muscles of G11-13 turtles and comparable stages of mouse are increasing volumetrically relative to body size (Fig. 3G,H).

**Fig 2.**
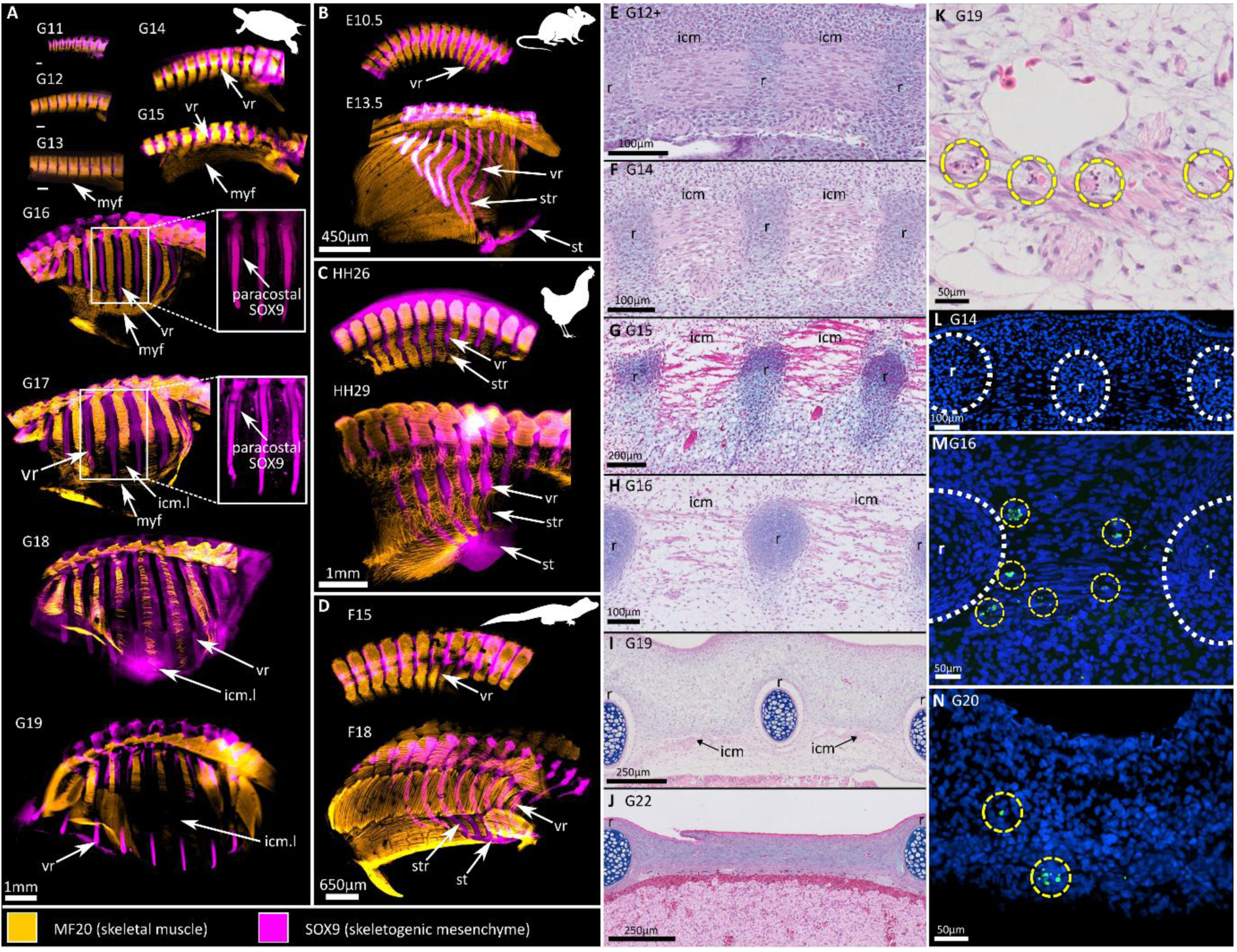
Comparative morphogenesis of the amniote body wall reveals turtle-unique dynamics. (A) Morphogenesis of the turtle body wall revealed by cleared, whole-mount embryos immunostained for early muscle (MF20) and cartilage (SOX9) markers. Early stages of rib and intercostal muscle development in turtles (G11-13) align closely with that of mouse (B), chicken (C), and alligator (D). These conserved early stages of rib and muscle development in turtle are confirmed by sagittally sectioned histology slides where highly organized skeletal muscle fills the intercostal space (E). This muscle begins to disorganize in the G14 turtle (F), a derived dynamic that is followed by a progressive loss of muscle fibres (G-J). Evidence of apoptotic activity in the intercostal spaces includes blebbing (K) and TUNEL assays (L-N) (see text for details). **Abbreviations:** icm, intercostal muscles; icm.l, intercostal muscle loss; st, sternum; str, sternal rib; vr, vertebral rib

**Fig 3.**
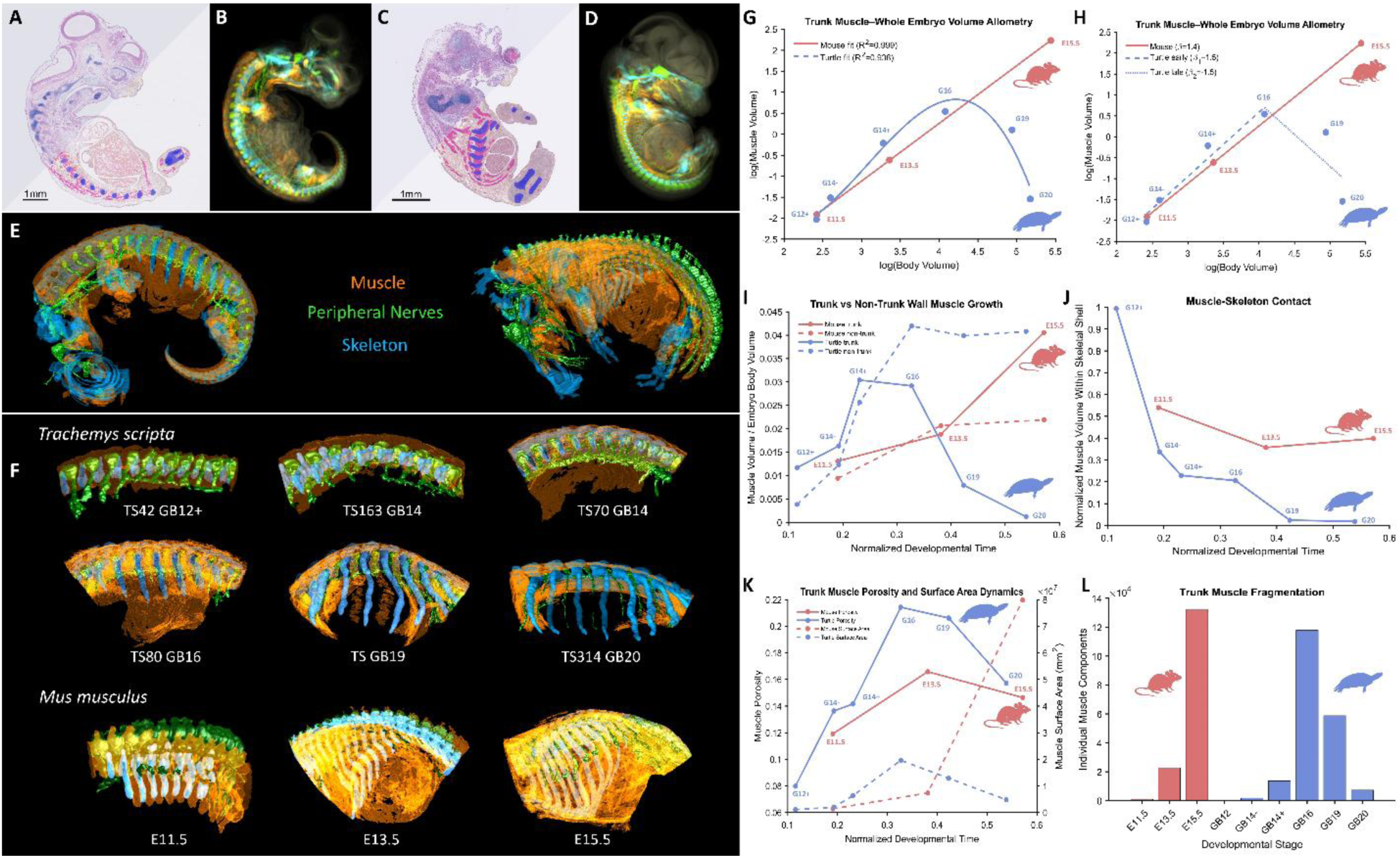
CODA-based quantitative 3D reconstruction reveals divergent muscle development in turtle and mouse embryos. (A-D) CODA-segmented tissues in representative thin sections shown alongside their 3D reconstructions in G16 turtle (A,B) and E13.5 mouse (C,D). Tissues include skeleton (cyan), muscle (orange), peripheral nerves (green), and a composite background of other tissues (e.g., viscera) (beige). (E) Whole-embryo 3D reconstructions of the same CODA-segmented tissues in G16 turtle (left) and E13.5 mouse (right). (F) Whole-embryo 3D reconstructions of the same CODA-segmented tissues in six stages of turtle and three stages of mouse; the reconstructions have been cropped between the pectoral and pelvic girdles to emphasize the region of interest. (G,H) Log–log scaling of muscle versus total embryo volume linear power-law scaling in mouse development. In contrast, turtle embryos exhibit nonlinear scaling, with early positive allometry up to G16 (β = 1.5) followed by negative scaling (β = −1.5). (I) Body wall muscle normalized to embryo volume shows sustained growth in mice, whereas turtles show an initial increase followed by a plateau and subsequent decline. Non–body wall muscle controls (e.g., limb and tail musculature) exhibit comparable growth between species. (J) Quantification of muscle–skeletal association reveals progressive detachment of musculature from the ribs in turtles, in contrast to stable muscle–rib connectivity in mice. (K,L) Muscle porosity, surface area dynamics, and fragmentation in the body wall illustrate initial muscular disorganization followed by progressive muscle loss in turtles.

The G14 embryo is where the turtle-unique features of the secondary body wall are first apparent. These features include the initial signatures of CR formation (ectodermal/mesenchymal thickenings), although CR maturation as a prominent, external ridge does not occur until G15 (Burke, 1989). The densely packed and craniocaudally aligned muscle fibres of the G14 myotome begin to disorganize (Figs. 2F, 3J). This disruption is in stark contrast to other amniotes whose intercostal myotome increases its organization, forming the tri-laminar morphology of external, internal, and innermost intercostal muscles whose orthogonal fibre orientation supports lung ventilation and thus respiratory function (e.g., Mekonen et al., 2015; Khabyuk et al., 2022). As with myogenesis (Buckingham, 2006), disorganization of the turtle myotome is first expressed in the most-cranial intercostal spaces. Signatures include loss of connectivity between adjacent muscle fibres/fascicles and a dissociation of muscle fibres from the ribs; by G16, this dissociation is complete (Fig. 2A). Histological and volumetric analyses further reveal increasing porosity, fragmentation, and thinning of muscle fibres, accompanied by an invasion of loose connective tissue that further disrupts the myofiber network (Figs. 2F-H; 3K,L). This structural disassembly may be realized via spindle-shaped, mesenchymal cells whose stellate-like projections extend towards muscle fibres and are most apparent in G16 (Supplementary Fig, 5A,B). These cells may be triggering apoptosis by inflicting some combination of structural and metabolic stress on the muscle fibres through the production of extracellular matrix and/or direct cell-cell or cell-matrix signalling. Regardless, the loss of mechanical and structural coupling that drives a transition from a continuous muscle sheet to discontinuous fragments, and is consistent with tissue-level atrophy and breakdown of organized contractile architecture (Mastaglia & Walton, 1971; Schiaffino et al., 2013). Notably, this structural disorganization occurs without an immediate reduction in total muscle volume. Between G14 and G16, trunk muscle remains a comparable proportion of total embryo volume (Fig. 3G,H), and early allometric scaling continues to follow the positive trajectory present in mouse (β ≈ 1.5 vs. β ≈ 1.4; Fig. 3H). The early steps in this derived phase thus include a profound disruption of tissue architecture, distinct from the conserved amniote program, rather than an immediate loss of muscle mass.

Volumetric analysis reveals G16 is coincident with a derived inflection in scaling behaviour where muscle growth in the turtle body wall transitions from positive to negative allometry (β ≈ −1.5; Fig. 3H). This shift corresponds to a subsequent decline in muscle volume normalized to embryo size that contrasts the sustained proportional growth in mouse (Figs. 2I; 3G-I). Importantly, growth in other muscle groups (e.g., limb and tail) remains comparable among species over these stages indicating turtle myogenesis is globally intact, with muscle loss restricted to the intercostal trunk (Fig. 3I). In contrast with myogenesis, ICM breakdown begins at the distal end of the intercostal space and proceeds proximally at a relatively slow rate. ICM fibres linger at the proximal end of this space for several stages but are absent by G22 (Fig. 2J) (hatching generally occurs at G26-27; Greenbaum, 2002).

Despite ICM degradation and loss, the turtle myotomal front and rib anlage continue to advance distally. Beginning at Stage 15, this advance occurs along different trajectories. The increasingly thick myotome includes an ancestral downward extension that enters the somatopleure and reaches the ventral body wall by G17. In contrast, the pre-chondrogenic ribs remain dorsal to the somatopleure by growing laterally. The rib tips penetrate the dorsal dermis and CR by G17, completing the turtle-unique process of axial arrest (Rice et al., 2015). Just prior to this penetration (G16), the paracostal gaps—created as ICM fibres disassociate from the ribs—become populated with a narrow field of Sox9 expression that is distinct from the rib core and absent in sampled outgroups (Fig. 1A-D). This expression field begins at the proximal end of the intercostal spaces and within the loose connective tissue of the subdermis. Subsequent expansion includes the entire intercostal breadth and nearly the entire length of the ribs. Its distal termination corresponds to the point of interface between rib and CR, beyond which the rib tip curves sharply downward. These intercostal tissues eventually lose Sox9 expression, presumably because, like the ribs, their maturation includes a collagenous framework of high osteogenic potential. By G25, this potential is realized through ossification of the dermal and subdermal layers of the intercostal tissues to form the T-shaped rib (Rice et al, 2015).

### Immune-Mediated Breakdown of Turtle ICM

Histological analysis of the intercostal spaces during ICM breakdown (G14-G22) reveals hallmarks of apoptosis, including nuclear condensation with hyperchromatic staining of polar bodies and blebbing (Aoki et al., 2020) (Fig. 2K; Supplementary Fig. 5). These features are absent at earlier stages and confirmed by TUNEL labelling of fragmented DNA (Gavrieli et al., 1992) (Fig. 2L-N). Apoptotic bodies persist through later stages, indicating sustained cell death with resultant cellular debris that must be cleared by phagocytic cells (Poon et al., 2014). Reptile immunology remains poorly characterized, limiting definitive identification of immune cells using specific markers (Field et al., 2022). Nevertheless, our histological data are consistent with localized innate immune activation accompanying developmental tissue remodelling. Phagocytic cells and macrophages become evident at G16 and persist through G20, in alignment with the onset and progression of ICM degradation. Increased vascularization is evident by G15, including blood-filled spaces, lined with a simple-squamous epithelium, that increase in both size and number during subsequent stages (Fig. 2K).

Importantly, this response does not resemble an acute inflammatory reaction as characterized by extensive heterophil/neutrophil invasion and tissue necrosis (Harmon, 1998; Kolaczkowska & Kubes, 2013). The pattern is more consistent with developmental or reparative inflammation associated with apoptotic tissue remodelling, including phagocytic recruitment, angiogenesis, extracellular matrix remodelling, and mesenchymal activation (Ribatti and Crivellato, 2009; Fantin et al., 2010; Stefater et al., 2011; Wood & Martin, 2017; Gordon & Plüddemann, 2018). The presence of eosinophil-like granulocytes may further support active remodelling and immune signaling through cytokine secretion, degranulation, and recruitment or modulation of macrophage populations (Abdala-Valencina et al., 2018). Although their role in reptilian embryogenesis remains poorly resolved, these cells could potentially contribute to extracellular matrix regulation or mesenchymal activation within the remodelling intercostal space.

The morphogenetic relevance of this inflammatory activity is unclear, but activated macrophages are known to secrete cytokines that promote angiogenesis, fibroblast activation, extracellular matrix remodeling, and osteogenic signaling, including BMP-associated pathways and TGF-β signaling (Sinder et al., 2015; Huang et al., 2021; Nacu et al., 2008). Non-pathological inflammation recently was implicated in an evolutionarily stable fusion of ancestral skeletal elements into a derived composite structure (i.e., pygostyle of birds; Rashid et al., 2023). Thus, the temporal coincidence of ICM loss, apoptosis, vascular invasion, innate immune recruitment, and subsequent intercostal ossification raises the possibility that inflammatory-remodelling pathways contribute directly to formation of the turtle-unique carapacial framework.

### Origin of the Turtle Body Plan – Synthesis and Extension

Here we used advanced 3D imaging of whole-embryo, amniote morphogenesis to promote our understanding of the developmental and evolutionary mechanisms underlying the origin of the turtle body plan. Our data define a biphasic developmental trajectory for the secondary body wall. The first phase is highly conserved and includes delamination and differentiation of an advancing sclerotome and myotome to produce a canonical set of thoracic, prechondrogenic ribs separated by an organized sheet of structurally integrated intercostal muscles. The second phase is characterized by a suite of turtle-unique transformations. These include well-documented CR formation and axial arrest, but also a novel disassembly of the ICM. This disassembly begins with disorganization, culminates with apoptotic regression and loss, and produces an overall growth trajectory that breaks the linear power-law scaling relationship expected for tissues forming adult structures. In this break, the trajectory aligns with the growth expectations of tissues serving critical early functions that diminish later in ontogeny. In metamorphic species, such tissues are abundant and function as part of a uniquely juvenile behavioural ecology; e.g., gills of aquatic tadpoles that regress in adult, terrestrial frogs (Etkin, 1932). In non-metamorphic species, these tissues are not as abundant and most often serve structural roles, such as patterning neighbouring tissues; e.g., notochordal patterning of the chordate neural tube (van Straaten et al., 1988).

Given the non-metamorphic status of turtles, we posit the turtle ICM develop as part of an inductive, morphogenetic system that includes rib patterning. An inductive role in rib formation and growth was recently promoted for ICM based on mouse and chick (Wood et al., 2020; Scaal, 2021; Khabyuk et al., 2022) and thus should be conserved across amniotes, including turtles. That advancement of the turtle myotome into the secondary wall precedes that of the rib-forming sclerotome meets the expectation of developmental primacy for tissues integrated under generative entrenchment (Wimsatt, 1986) and aligns with the developmental sequence also observed in mouse, chick, and alligator (Scaal, 2021; Khabyuk et al., 2022; our data). The G14 correlation between initiation of ICM disassembly and lateralization of the ribs (axial arrest) constitutes further evidence that these two tissues are developmentally and not just structurally linked. In short, turtle ICM continue to differentiate because they are developmentally integral to the ribs, which are structurally integral to the shell and thus organismal viability.

The G14 embryo also contains the first evidence of CR formation. It is unclear whether the CR signalling hypothesized to drive axial arrest (Burke, 1989; Loredo et al., 2001; Cebra-Thomas et al., 2005) would be in place during this early organizational phase or whether its phenotypic efficacy is realized only with CR maturity at G15. There is currently no evidence that the CR is in an instructive relationship with the ribs at G14; rather, its earliest molecular activity relates to internal proliferation and organization (Moustakas-Verho, 2008; Zhang et al., 2021). If CR effects begin only at G15, then ICM disassembly enjoys clear developmental primacy and is unlikely to be under CR control. In this scenario, ICM breakdown moves to a foundational position within the turtle-unique morphogenesis of the body wall, with CR playing the later, but still critical, role of inducing axial arrest. If CR signalling, however, is coincident with the first signatures of CR structural organization, then it remains possible the CR patterns all turtle-unique aspects of the biphasic secondary body wall, including ICM breakdown.

Such wide-ranging effects align with the evolutionary hypothesis that the CR represents a macromutational innovation capable of producing the modern turtle body plan in a geological instant, with no expectation of a transitional fossil record. The problem for this ‘hopeful monster’ scenario is that a transitional fossil record not only exists for the turtle body plan but establishes ICM regression and costal formation (expression of the T-shaped rib) as preceding axial arrest by upwards of 40my; the c.260my *Eunotosaurus africanus* and c.220my *Odontochelys semitestacea* being the respective early and late constraints on this inferred interval. Removing *Eunotosaurus* from the turtle stem lineage (Simões et al., 2022) may reduce this interval by as much as half, but otherwise makes no other requirements of the evo-devo model presented here. The requirement that is imposed by the removal of *Eunotosaurus* is that the dramatic early stages of trunk repatterning described here and elsewhere—including the acquisition of a highly derived abdominal sling that drives lung ventilation (Lyson et al. 2016)—evolved at least twice.

Given these data, the correlation between ICM degeneration and rib transformation is likely causative. Loss of this muscle-based signalling centre would have liberated the later-stage ribs from the constraints of their ancestral patterning, providing them with renewed morphogenetic potential (Fig. 4A,B). Some of this potential remained latent until the CR evolved and engaged the developing ribs in a novel signalling relationship that resulted in their axial arrest (Fig. 4C). Other potential was realized with more immediacy, possibly flowing directly from the inflammatory response that mediates ICM breakdown and is known to initiate BMP signalling capable of producing metaplastic bone (Sinder et al., 2015; Huang et al., 2021) (Fig. 4D). A novel observation integral to this model is the paracostal band of subdermal Sox9 expression that expands in the wake of muscle degeneration to fill the intercostal space. This bridge of intercostal tissue bearing high osteogenic potential constitutes the structural base on which subsequent dermal ossification forms the T-shaped ribs of both perinatal modern turtles and early turtle-line fossils and serves as the structural framework for the mature carapace. That the inflammation associated with muscle breakdown initiates: 1) the paracostal gap between ICM and rib, 2) the Sox9 expression within this gap, 3) the intercostal expansion of this expression as the muscle is cleared, or 4) some combination of the three, are all possibilities congruent with our data.

**Fig 4.**
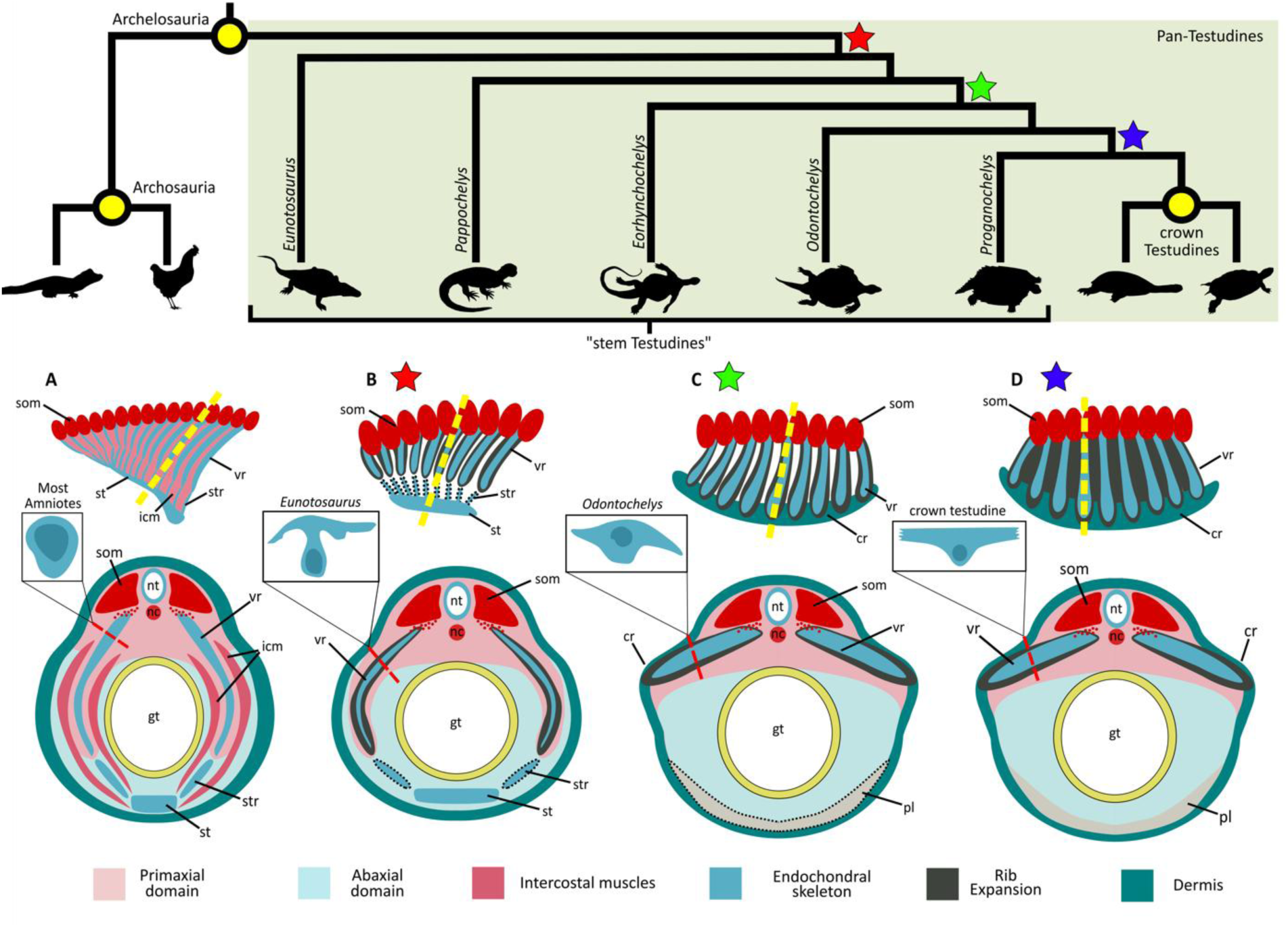
Expanded evolutionary hypothesis for the origin of the turtle carapace incorporating novel developmental and fossil-based observations. (A) The base of the archelosaur radiation was characterized by a conserved thoracic body wall wherein an organized intercostal musculature (ICM) directs the developing ribs into the ventral body wall. (B) Early in the turtle stem lineage, the trunk musculature underwent profound transformations with the products being an alternative mechanism for lung ventilation followed by late-stage loss of the ICM. The dynamics responsible for ICM loss also initiated a derived ossification of the intercostal dermis, producing uniquely widened ribs that retain their ancestral orientation (as found in *Eunotosaurus* and *Pappochelys*). (C) Axial arrest of the ribs (as found in *Eorhynchochelys, Odontochelys*, and all crownward turtles). This rib lateralization likely marks the appearance of the carapacial ridge, or at least its ability to draw the ribs out of their plesiomorphic trajectory following ICM loss. (D) Completion of the modern turtle carapace (as found in *Proganochelys* and crownward turtles). Yellow dotted lines indicate position of accompanied coronal sections. Red dotted lines show location of rib cross section. Black dotted lines indicate structures present in some, but not all taxa included in each phase. Cross sections of rib shafts redrawn from Lyson et al. (2014) and Schoch and Sues (2015). **Abbreviations:** cr, carapacial ridge; gt, gut tube; nc, notochord; nt, neural tube; pl, plastron; som, somites; st, sternum; str, sternal rib; vr, vertebral rib.

Support for a mechanism that can produce the T-shaped rib of turtles in lieu of axial arrest aligns the developmental and paleontological signatures of the turtle body plan and its initiation. This new model also clarifies why the paracostal gap, paracostal Sox9 expression, and finally the mature, T-shaped morphology of the rib occur only proximal to the point where the ribs penetrate the dermis. The co-ossification of ribs and dermis does not require ensnarement of the former by the later. In fact, this ensnarement actually negates localized co-ossification, probably by disrupting the unique subdermal morphogenesis that serves as its developmental and structural precursor. Muscle loss as a critical mechanism in the development and evolution of a derived character system is known in other amniotes (Tran et al., 2019; Tran et al., 2024). Only in turtles, however, do we find such profound, embryonic loss, with morphogenetic effects that rise to the level of a novel body plan. What causative dynamics lie upstream of ICM disorganization and loss remains unclear. If the ribs are the source of the stellate cells implicated in myogenic disruption and apoptosis initiation, then the rib and ICM are engaged in a successive series of inductions and disruptions that first promoted intercostal ossification and set the stage for subsequent axial arrest.

More than 250 million-years ago, a relatively young lineage of sauropsid reptiles evolved the seemingly unremarkable variation of T-shaped ribs (Lyson et al., 2013; Schoch & Sues, 2015; Lyson and Bever, 2020). This novel costal morphology stiffened the trunk, conveying mechanical advantage to the limbs and thus selective benefits for either digging or swimming—two behaviours that characterize a large proportion of subsequent turtle diversity and ecology (Lyson et al., 2014; Lyson et al., 2016). From this seemingly inauspicious beginning, the turtle lineage realized what assuredly is the most extreme known example of hyperossification in the history of terrestrial vertebrates. That stated, it is increasingly clear that the story of turtle origins is written as much in the muscles as it is in the bones. It was a transformation of the abdominal muscles, producing a redundancy in lung ventilation, that allowed the trunk to initially stiffen and ossify (Lyson et al., 2016). Folding of the shoulder muscles granted the modern shell its status as a viable pathway (Nagashima et al., 2009; Kuratani et al., 2011). Here, we add to this trend with support for ICM disassembly as a critical bridge in this transformational history, one that will hopefully catalyse a new research cycle on this remarkable natural experiment.

## SUPPLEMENTARY INFORMATION

### Animal Husbandry

Eggs, and in the case of mice, embryos, were sourced from state-registered wildlife refuges, commercial breeders, and scientific suppliers (Supplementary Table 1). Egg incubation, tissue extraction, and paraffin-based histology workflows followed methodologies outlined in Gensbigler et al. (2025). Eggs were incubated under species-specific conditions, including temperature, humidity, and substrate, as established in the literature (Hamburger & Hamilton, 1951; Ferguson, 1985; Marlen & Fischer, 1999; Greenbaum, 2002; Tulenko & Sheil 2007).

Embryos were euthanized via metabolic suppression in a 4°C freezer and extracted using species-tailored dissection protocols (see Gensbigler et al., 2025). Late-stage embryos were surgically decapitated following best practices outlined by the Institute for Laboratory Animal Research (Institute for Laboratory Animal Research, 2010) and the American Veterinary Medical Association (Underwood & Anthony, 2020). Embryonic stages for all taxa were assigned based on published criteria (Supplementary Table 1). Following extraction, amniotic membranes were removed, and embryos were rinsed in 1X phosphate buffered saline (PBS) before fixation in neutral buffered formalin for durations dependent on embryonic size (see Gensbigler et al., 2025). Embryos were dehydrated in ethanol for classical histology or methanol for CLARITY protocol (Supplementary Table 2).

**Supplementary Table 1.**
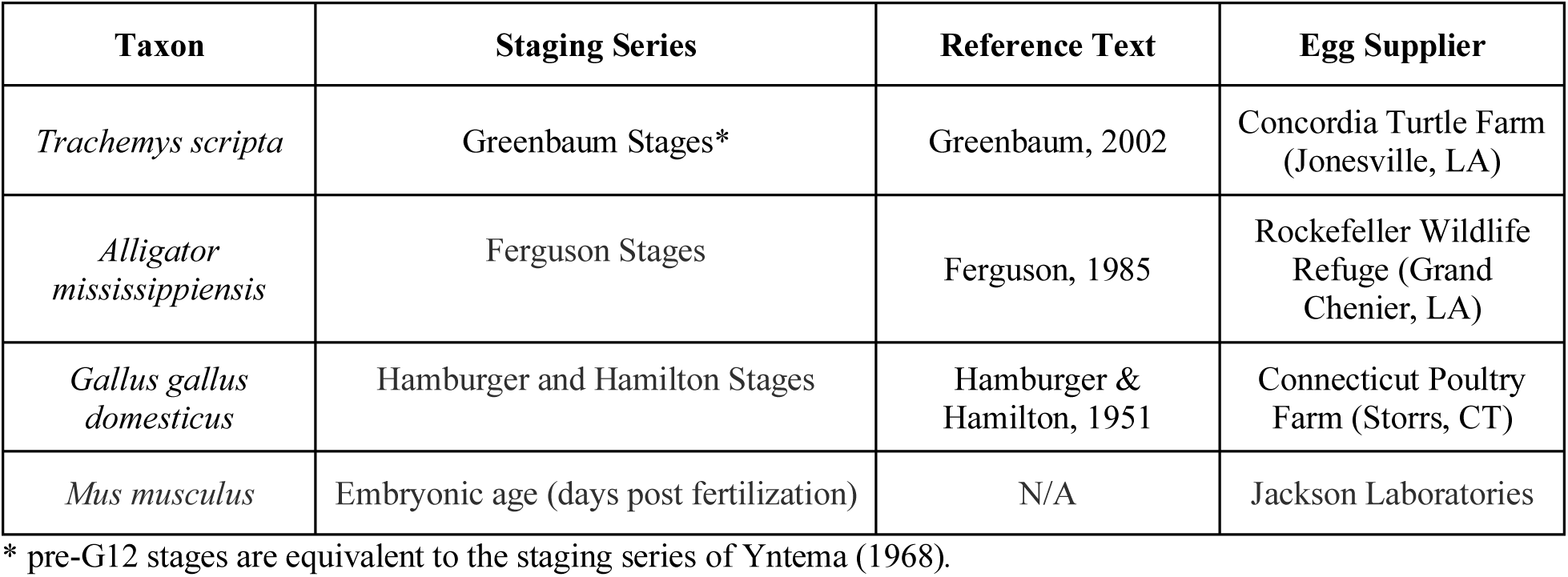
Taxon-specific staging series for comparative embryogenesis.

**Supplementary Table 2.** Embryos and their usage in this study.

| **Taxon** | **Stage** | **Days Post Fertilization** | **Application** |
| --- | --- | --- | --- |
| *Mus musculus* | N/A | 11.5 | 3D reconstruction and quantification |
| *Mus musculus* | N/A | 13.5 | 3D reconstruction and quantification |
| *Mus musculus* | N/A | 15.5 | 3D reconstruction and quantification |
| *Trachemys scripta* | G12+ | 14 | 3D reconstruction and quantification |
| *Trachemys scripta* | G14- | 18 | 3D reconstruction and quantification |
| *Trachemys scripta* | G14+ | 20 | 3D reconstruction and quantification |
| *Trachemys scripta* | G16 | 25 | 3D reconstruction and quantification |
| *Trachemys scripta* | G19 | 30 | 3D reconstruction and quantification |
| *Trachemys scripta* | G20 | 36 | 3D reconstruction and quantification |
| *Mus musculus* | N/A | 10.5 | CLARITY |
| *Mus musculus* | N/A | 13.5 | CLARITY |
| *Gallus gallus* | HH24 | 4 | CLARITY |
| *Gallus gallus* | HH29 | 6.5 | CLARITY |
| *Alligator mississippiensis* | F15 | 18-20 | CLARITY |
| *Alligator mississippiensis* | F18 | 24-26 | CLARITY |
| *Trachemys scripta* | Y11 | 10-14 | CLARITY |
| *Trachemys scripta* | G12 | 14-16 | CLARITY |
| *Trachemys scripta* | G13 | 16-19 | CLARITY |
| *Trachemys scripta* | G14 | 19 | CLARITY |
| *Trachemys scripta* | G15 | 21 | CLARITY |
| *Trachemys scripta* | G16 | 25 | CLARITY |
| *Trachemys scripta* | G17 | 28 | CLARITY |
| *Trachemys scripta* | G18 | 29 | CLARITY |
| *Trachemys scripta* | G19 | 32 | CLARITY |
| *Trachemys scripta* | G15 | 19 | Histology |
| *Trachemys scripta* | G18 | 27 | Histology |
| *Trachemys scripta* | G22 | 42 | Histology |

### Wholemount Tissue Clearing of Amniote Embryos

The modified CLARITY (Chung et al., 2013) workflow used in this study is largely based on the original protocol of Griffin et al. (2022), but includes modifications recommended by Matteo Fabbri (personal communication). All steps requiring an ambient temperature of 4℃ took place within a commercial refrigerator. Steps at 37℃ occurred within a heated water bath. Unless otherwise noted, all other steps were carried out at room temperature. Specimens were kept in glass test tubes throughout the workflow, and all effort was made to reduce the number of transfers between containers, as this is when embryos are most likely to be damaged. To reduce the risk of inadvertent damage to the specimens, transfer pipettes were used when swapping out solutions, and large volumes of liquid were never poured or squirted directly onto embryonic tissue, especially at early developmental stages.

Formalin-fixed embryos were gradually dehydrated via four 15-minute washes in methanol-1X PBS (phosphate buffered saline) solutions of incrementally decreasing PBS concentration (¼ methanol-1X PBS, ½ methanol-1X PBS, ¾ methanol-1X PBS, and 100% methanol). For all washes, samples were immersed in a total volume roughly ten times that of the embryo to ensure complete tissue permeation. Embryos were then bleached in an 8:1:1 solution of Methanol, DMSO, and 30% hydrogen peroxide. Bleaching removes melanin from pigmented tissues, promoting subsequent clearing procedures. Embryos were bleached for at least 24 hours, but later stages, particularly those of *Mus* and *Trachemys*, required up to 30 hours. We found that yellowing of the eyes was a good external indicator that the embryo had been sufficiently bleached. Following the bleaching step, embryos were completely rehydrated in four 15-minute washes of reversed concentrations of the initial dehydration step (¼ 1X PBS-Methanol, 2/4 1X PBS-Methanol, ¾ 1X PBS-Methanol, and 100% 1X PBS). Embryos in 100% 1X PBS were stored at 4℃ for several days.

Samples were next submerged in a hydrogel solution that invests embryonic tissues with an antibody-permeable gel whilst maintaining tissue integrity. A hydrogel solution was mixed in 200ml batches consisting of 160ml de-ionized water, 20ml of 10X PBS, and 20ml of 40%. Immediately prior to embryo submersion, 0.5g of VA-044 was added as polymerization initiator. Embryos were left under motion overnight at 4℃.

To trigger the hydrogel monomer reaction, embryo tubes without lids/caps were mounted in a test tube rack and placed into a 37℃ water bath under motion for three-to-four hours. To provide motion, the bath was placed onto a wheeled cart, and a nutating mixer set to a high RPM was placed on a lower shelf, causing the whole cart to oscillate slightly. The water level in the bath was high enough to submerge the test tubes to approximately mid-height. Observation of oxygen bubbles within the solution after two-to-three hours indicated that the reaction had begun. The hydrogel solution was removed from each tube after four hours and replaced with 1XPBS. The tubes were capped and returned to the water bath for one hour. This wash was repeated five more times, and samples were left at 4℃ overnight in fresh 1XPBS. The following day, samples were washed five times under motion and at 37℃ in a PBSt solution (1XPBS, 0.5% Triton) for one hour each wash. Triton is a surfactant that improves antibody permeation by reducing the effects of surface tension between solutions. Samples were left in fresh PBSt overnight at 4℃, after which they were ready for wholemount immunohistochemistry.

### Wholemount Immunohistochemical Staining of Amniote Embryos

To observe the morphogenetic activity of trunk tissues, indirect immunofluorescence was applied to tag three proteins of interest in each embryo: SOX9 (endochondral skeletal precursors), MF20 (skeletal muscle), and HNK-1 (neural crest derivatives) (data not shown). Concentrations of each antibody were optimized to maximize tissue coverage and minimize non-specific binding (Supplementary Table 3). Each primary antibody was targeted to a different host or isoform to prevent cross-reactivity between fluorescence channels. Primary antibody solutions were mixed in 10ml batches composed of PBSt (8,590µl), DMSO (500µl), horse serum (500µl), 0.5% sodium azide (100µl), SOX9 (10µl), MF20 (250µl), and HNK-1 (50µl). Enough primary solution was added to the tubes so that the embryos were totally submerged. Tubes were left in a 37℃-water bath under motion for four days.

After four days, the primary antibody solution was replaced with PBSt and washed five times at 37℃ under motion. Fluorescent secondary antibodies were selected to minimize spectral overlap and to bond exclusively to each primary antibody (Supplementary Table 4). Secondary antibody solutions were mixed in 10ml batches composed of PBSt (9300µl), DMSO (500µl), 0.5% sodium azide (100µl), anti-SOX9 (33.3µl), anti-MF20 (33.3µl), and anti-HNK-1 (33.3µl). Embryos were left in secondaries at 37℃ under motion for four days. To prevent fluorophore photobleaching, the water bath was covered with a foil sheet to maintain low-light conditions. After four days, embryos were washed four times in PBSt under motion at 37℃.

### Fluorescent Microscopy and Image Capture

Immunostained stained embryos were prepared for imaging through equilibration in a refractive index matching solution (RIMS). This solution was made in batches composed of 30ml 0.002 M phosphate buffer, 30µl of 10% sodium azide, and 40g of Histodenz non-ionic density gradient medium (Millipore Sigma D2158). To fully dissolve the Histodenz powder, the solution was left in an incubator oven set to 40℃ and stirred occasionally over three hours. Embryos were equilibrated in RIMS for two days at 4℃. Samples were be stored in RIMS at 4℃ for multiple months (recommendation: leave specimens in the back of a fridge in a box to reduce temperature and light-level perturbations).

Immediately prior to confocal imaging, embryos were mounted in glass-bottomed observation dishes (CellTreat Scientific Products 229632). To prevent the sample moving during z-stack acquisition, excess RIMS was removed with a transfer pipette so that the embryo settled on the bottom of the dish. Embryos were gently positioned using a fine paintbrush, or with the tip of a plastic transfer pipette. Prior to imaging via light sheet microscopy, RIMS was removed from specimen tubes and replaced with glycerol roughly two hours prior to imaging. The sample chamber was also filled with glycerol. Embryos were secured to the observation stage using a small drop of super glue and slowly lowered into the filled chamber. Following light sheet imaging, embryos were removed from the instrument by gently breaking the super glue bond with the observation stage. Specimens were then returned to RIMS solution and stored at 4℃. Observations made during these experiments suggest that specimens may be successfully mounted and imaged up to three times prior to total photobleaching or critical tissue damage.

**Supplementary Table 3.** Immunohistochemical targets, concentrations, and supplier of primary antibodies.

| Protein | Target Tissues | Host and Isotype | Concentration | Supplier |
| --- | --- | --- | --- | --- |
| SOX9 | Early endochondral precursors and chondrocytes | Rabbit (Na) | 1:1000 | Sigma Aldrich (AB5535) |
| MF20 | Skeletal muscle | Mouse (IgG2b) | 1:40 | DSHB (MF20) |
| HNK-1 | Neural crest derivatives | Mouse (IgG1) | 1:300 | DSHB (1C10) |

**Supplementary Table 4.** Host species, isotype target, concentration, and supplier of secondary antibodies.

| Channel | Host/Isotype | Host/Isotype Target | Concentration | Supplier |
| --- | --- | --- | --- | --- |
| Channel 1 (SOX9) | Donkey | Anti-Rabbit | 1:300 | ThermoFisher (A-31573) |
| Channel 2 (MF20) | Goat | Anti-Mouse (IgG2b) | 1:300 | ThermoFisher (A-21147) |
| Channel 3 (HNK-1) | Goat | Anti-Mouse (IgG1) | 1:300 | ThermoFisher (A-21121) |

### Classical Histology

Fresh embryos were formalin-fixed and paraffin-embedded (FFPE) using a Leica ASP300S fully enclosed automated vacuum tissue processor. Processing runs were programmed using size-dependent dehydration, xylene clearing, and wax infiltration periods as described in Gensbigler et al. (2025). Thin sections were cut from embedded samples using a Leica Histocore Biocut microtome. Specimens were exhaustively serially sectioned in sagittal plane at 4 µm thickness and mounted on Rankin charged microscope slides (product no. 20245W). Two serial sections were placed on the same slide to minimize the number of slides used. For the largest specimen (TS314), approximately two-thirds of the embryo was sectioned due to size constraints; one-half of the specimen was used for downstream quantification and visualization. Tissues were stained using Alcian Blue Hematoxylin/Orange G Eosin (H&E and alcian) staining methods (see Gensbigler et al., 2025; Table 6) using a Leica ST5020 automated slide stainer, and coverslips were applied using a Leica CV5030 automated cover slipper. Slides were imaged at 20x magnification for 3D reconstruction or 40x magnification for histological visualization using a Hamamatsu NanoZoomer S360 brightfield slide scanner.

### TUNEL Assay

TUNEL (Terminal deoxynucleotidyl transferase dUTP Nick End Labeling) was performed using the ApopTag® Fluorescein In Situ Apoptosis Detection Kit (Millipore Sigma product no. S7110) following the manufacturers protocol. Paraffin embedded embryos were sectioned in sagittal plane at 6 µm and mounted on Superfrost Plus TM slides. Sections were deparaffinized in xylene (3 × 5 min washes), rehydrated through a graded ethanol series, and rinsed in 1X PBS. Sections were then permeabilized using proteinase K digestion. Terminal deoxynucleotidyl transferase (TdT) was used to incorporate digoxigenin-labeled nucleotides at sites of DNA fragmentation. Labeled DNA was detected using an anti-digoxigenin fluorescein-conjugated antibody. Slides were mounted using VECTASHIELD® HardSet™ Antifade Mounting Medium with DAPI and stored at −20°C away from a light source. Fluorescent images were acquired using a Nikon Eclipse Ti2-E with Yokogawa CSU-W1 SoRA spinning disk confocal microscope.

### CODA 3D modeling

#### Image Pre-Processing

Whole-slide images (WSIs) of stained tissue sections acquired in NDPI format at 20× magnification (0.5 µm per pixel) were processed using OpenSlide to generate downsampled TIFF images at pseudo-1x (10 µm per pixel), pseudo-5x (2 µm per pixel), and pseudo-10x (1 µm per pixel) magnification for computational analysis. As most images contained multiple tissue sections, downsampled images were cropped using software written in MATLAB 2024b to generate individual TIFF images and coordinates containing a singular tissue section per file. Cropped pseudo 1x, 5x, and 10x images were used for downstream image registration, semantic segmentation, and 3D rendering tasks.

#### Nonlinear Image Registration

Cropped image stacks were aligned to reconstruct 3D volumes using a modified version of previously described methods (Kiemen et al., 2020). Specifically, they were adapted to accommodate the non-uniform geometry of embryonic tissues. Sections exhibiting substantial sectioning or staining artefacts were excluded prior to alignment to prevent propagation of registration artifacts. Of the remaining high-quality sections in each sample, the center image was assigned as the reference coordinate system. Alignment of each image into the reference coordinate system was calculated iteratively on image pairs in two stages. First, rigid registration was used determine the slide-level rotation and translation required to best globally align adjacent slides. Second, nonlinear registration was used to determine local adjustments necessary to correct for tissue shrinkage or expansion, tearing, folding, and other local artefacts. Elastic registration was implemented using a tile-based approach, where the tile size increased progressively with distance from the embryonic midline. This adaptive strategy minimized over-warping in peripheral regions while preserving the native fusiform morphology of the embryo, which exhibits progressive tapering away from the central body axis.

#### Semantic Segmentation of Tissue Types from Histology

Deep learning-based semantic segmentation models were trained to label desired morphology in the embryonic histology images using the CODAvision software (Matos et al., 2025). For each specimen, a resNet50 model modified for semantic segmentation using the deeplabV3 plus workflow (Chen et al., 2017) was finetuned to segment the desired histological structures.

A set of histologically recognizable tissue types were defined to exhaustively classify different anatomical regions in the embryos (Supplementary Table 5). For each embryo, a subset of 15-20 histology images sampled across the image stack were extracted. On each image, the examples of each tissue type present were manually annotated using Aperio ImageScope. For each model, a total of 30 to 200 examples of each tissue class were generated. An independent set of annotations was generated to test the models on images unseen during model training. The precision of each model was assessed, and the training annotations were refined until the per-class precision and recall exceeded 85% (Sup. Fig 1). The final models were applied to the full cohorts of downsampled, cropped pseudo-10x images (1 µm per pixel) to exhaustively label desired anatomical structures in the histology.

**Supplementary Table 5.** Standardized tissue classes across mouse and turtle developmental stages.

| Tissue Class | Mouse E11.5 | Mouse E13.5 | Mouse E15.5 | Turtle (G12) | Turtle G14 | Turtle G14 | Turtle G16 | Turtle G19 | Turtle G20 |
| --- | --- | --- | --- | --- | --- | --- | --- | --- | --- |
| Extracellular Matrix | ECM | ECM | ECM | ECM | ECM | ECM | ECM | ECM | ECM |
| Blood / Vasculature | Blood/Vasc | Blood/Vasc | Blood/Vasc | Blood/Vasc | Blood/Vasc | Blood/Vasc | Blood/Vasc | Blood/Vasc | Blood/Vasc |
| Heart | Heart | Heart | Heart | Heart | Heart | Heart | Heart | Heart | Heart |
| Eye | Eye | Eye | — | Eye | Eye | Eye | Eye | Eye | — |
| Cartilage | Cartilage | Cartilage | Cartilage | Cartilage | Cartilage | Cartilage | Cartilage | Cartilage | Cartilage |
| Central Nervous System | CNS | CNS | CNS | CNS | CNS | CNS | CNS | CNS | CNS |
| Peripheral Nerves | Nerves | Nerves | Nerves | Nerves | Nerves | Nerves | Nerves | Nerves | Nerves |
| Muscle | Muscle | Muscle | Muscle | Muscle | Muscle | Muscle | Muscle | Muscle | Muscle |
| Lungs | — | Lungs | Lungs | — | — | — | — | Lungs | Lungs |
| Liver | Liver | Liver | Liver | — | — | — | — | Liver | Liver |
| Kidney | Kidney | Kidney | Kidney | Kidney | Kidney | Kidney | Kidney | Kidney | Kidney |
| Mesonephros | — | — | — | Mesonephros | — | Mesonephros | Mesonephros | Mesonephros | Mesonephros |
| Skin | Skin | Skin | Skin | Skin | Skin | Skin | Skin | Skin | Skin |
| Epithelium | Epithelium | Epithelium | Epithelium | Epithelium | Epithelium | Epithelium | Epithelium | Epithelium | Epithelium |
| Bone | Bone | — | Bone | — | — | Bone | Bone | — | Bone |
| Ribs | Ribs | Ribs | Ribs | — | Ribs | Ribs | Ribs | Ribs | — |
| Gland | Gland | — | Gland | Gland | — | — | — | — | Gland |
| Esophagus | — | — | Esophagus | — | — | — | — | — | — |
| Thymus | — | — | Thymus | — | — | — | — | — | — |
| Trachea | — | — | Trachea | — | — | — | Trachea | — | — |
| Other Internal Organs | Other | Other | Other | Other | Other | Other | Other | Other | Other |

**Supplementary Fig 1.**
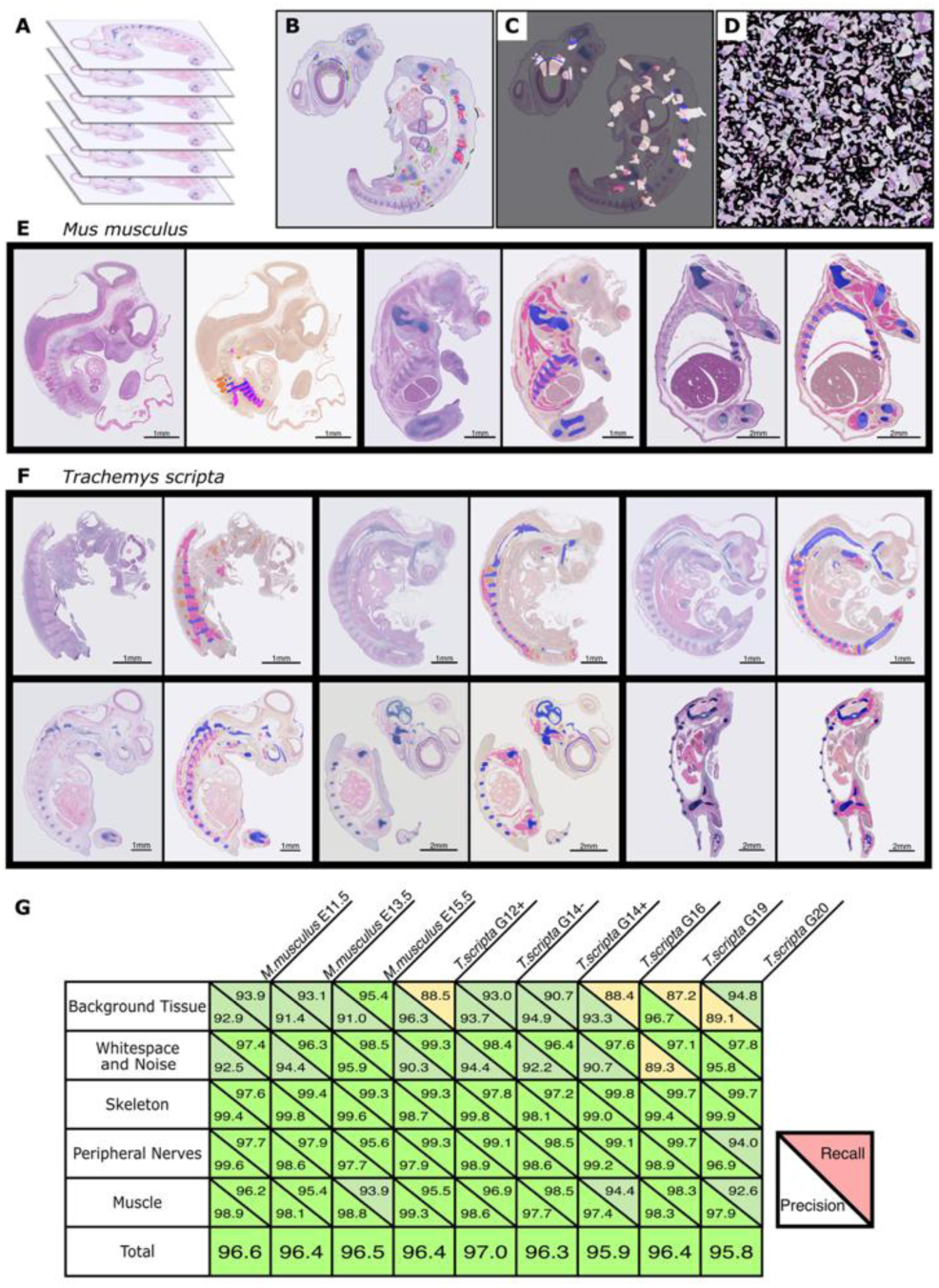
CODA-based 3D reconstruction, annotation, and segmentation performance. (A) For 3D reconstruction, embryos were paraffin embedded, exhaustively serially sectioned at 4 µm, stained, and imaged at 0.5 µm per pixel. (B) A subset of digitized sections was annotated and used to train a segmentation algorithm to fully segment whole slide images seen in (C). (D) Data augmentation using big-tile strategy, in which annotated regions are extracted and systematically tiled to generate training patches while preserving local spatial context. Representative H&E and alcian sections alongside corresponding CODAVision semantic segmentations for mouse (E) and turtle (F) developmental series, with muscle (pink), skeletal tissue (blue), nerves (orange), and background (beige), used for downstream visualization and quantification. (G) Model performance across all specimens showing per-tissue class precision (bottom) and recall (top), as well as overall accuracy; all tissue classes exceed 85% accuracy, with all overall model accuracy exceeding 95%.

#### Region of Interest Generation

To isolate relevant thoracic and abdominal trunk anatomy for quantification, a region of interest (ROI) was defined on semantically segmented image stacks using MATLAB Volume Segmenter (Sup Fig 2). To ensure anatomical consistency across species despite differences in the axial skeleton the ROI was bounded using conserved skeletal landmarks. Specifically, the pectoral and pelvic girdles were used to define the anterior and posterior limits of the trunk region, respectively. This landmark-based approach enabled consistent extraction of homologous anatomical regions across all specimens. Segmented image stacks were cropped to this ROI, retaining only tissues within the trunk domain for all downstream analyses and visualization. A secondary “body” ROI was defined for whole-body measurements. In this case, the head region was excluded to ensure consistency across specimens, as later-stage embryos were decapitated in accordance with ethical guidelines. This standardized body ROI enabled direct comparison of tissue volumes and proportions across developmental stages and species.

**Supplementary Fig 2.**
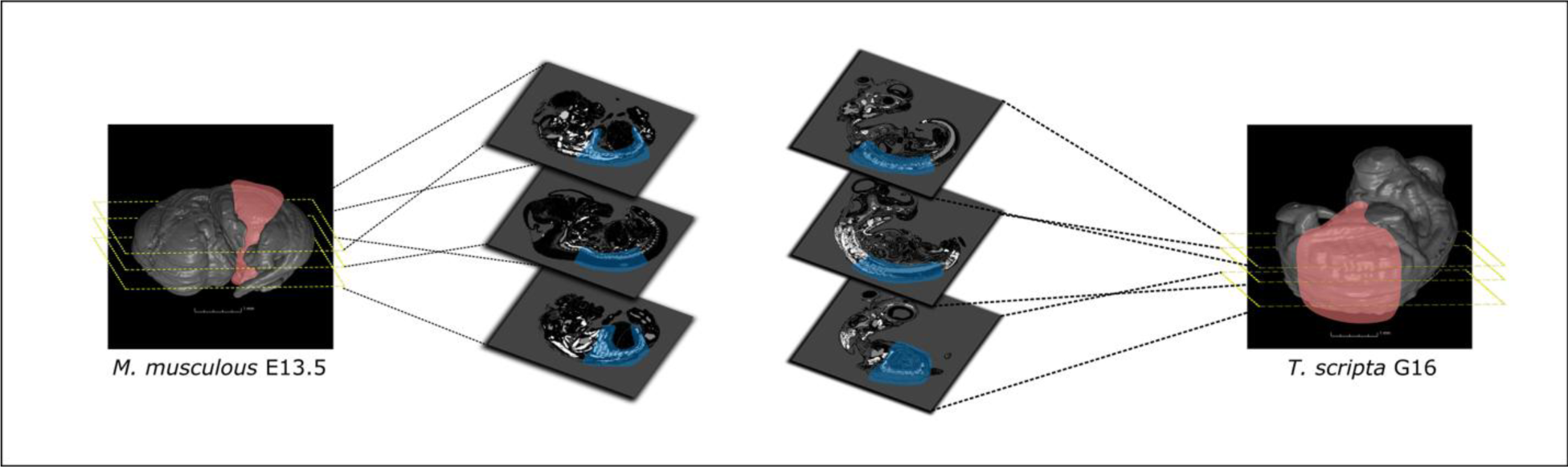
Region of interest generation. Region of interest generation using MATLAB Volume Segmenter (mouse, left; turtle, right). Aligned CODA 2D semantic segmentation outputs were manually cropped and interpolated to isolate the trunk region between the pectoral and pelvic girdles. These cropped slices were then reconstructed to generate the 3D ROIs (red region) used for downstream quantitative analyses.

#### Three-Dimensional Tissue Visualization

Segmented masks were aligned to a common coordinate space through application of the registration transforms calculated in the pseudo-1x H&E and alcian space. Registered, segmented masks were loaded into MATLAB 2024b, downsampled four-fold such that the axial resolution matched the lateral resolution (corresponding to the tissue thickness of 4µm) and compiled into a three-dimensional matrix. In the matrix, voxels contained pixel values corresponding to the tissue classes labelled using the segmentation models and possessed isometric dimensions of 4×4×4 µm^3^. These matrices were used for all downstream three-dimensional histology visualization and quantification tasks.

Mesh generation and rendering were performed in MATLAB using the isosurface and patch functions. Visualization was restricted to selected tissue classes: muscle, peripheral nerve, and skeleton, to emphasize structures relevant to musculoskeletal morphogenesis and reduce visual occlusion from unrelated tissues.

Meshes were exported in STL format and processed in MeshLab (v2025.07). To improve visualization of trunk anatomy, ROI volumes were bisected along the sagittal plane, enabling unobstructed viewing of structures and tissues. Small, disconnected components arising from segmentation noise were removed. To mitigate z-depth stair-step artifacts introduced by the sectioning thickness, three iterations of HC Laplacian smoothing were applied to skeletal and neural meshes. Final STL meshes were imported into VG Studio (2025) for rendering. Tissue types were assigned surface colors to enhance perception of depth and spatial interpretation (Sup Fig 3). Images for use in figures were exported directly from VG Studio.

#### Quantification

Raw, unmodified outputs from the segmentation algorithm restricted to the region of interest (ROI) corresponding to the trunk body wall between the girdle systems were used for all analyses. All quantification and data visualization were performed in MATLAB 2024b.

**Supplementary Fig 3.**
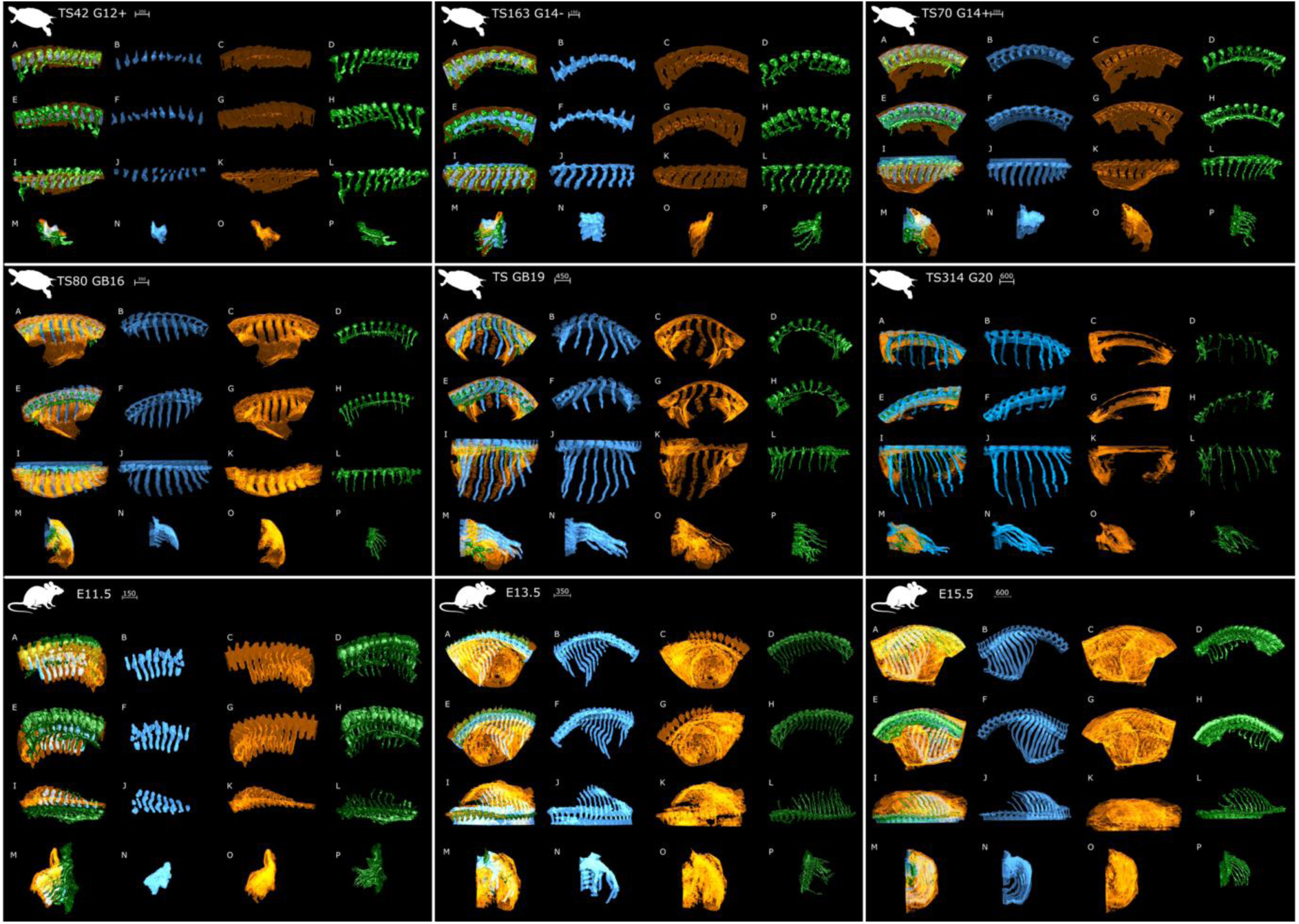
3D reconstructions of trunk tissues across developmental stages in turtle and mouse embryos. Each panel shows a reconstructed embryo within the trunk region of interest. The top six panels correspond to *Trachemys scripta* embryos, and the bottom three to *Mus musculus*. Within each panel, columns display (left to right): (1) sagittally bisected embryo with reconstructions of muscle (orange), skeleton (blue), and peripheral nerves (green); (2) isolated skeleton; (3) isolated muscle; and (4) isolated peripheral nerves. Rows correspond to different viewing orientations: lateral view of the left hemi-embryo (top row), lateral view of the right hemi-embryo (second row), dorsal view of the left hemi-embryo (third row), and anterior view of the left hemi-embryo (bottom row).

**Supplementary Fig 4.**
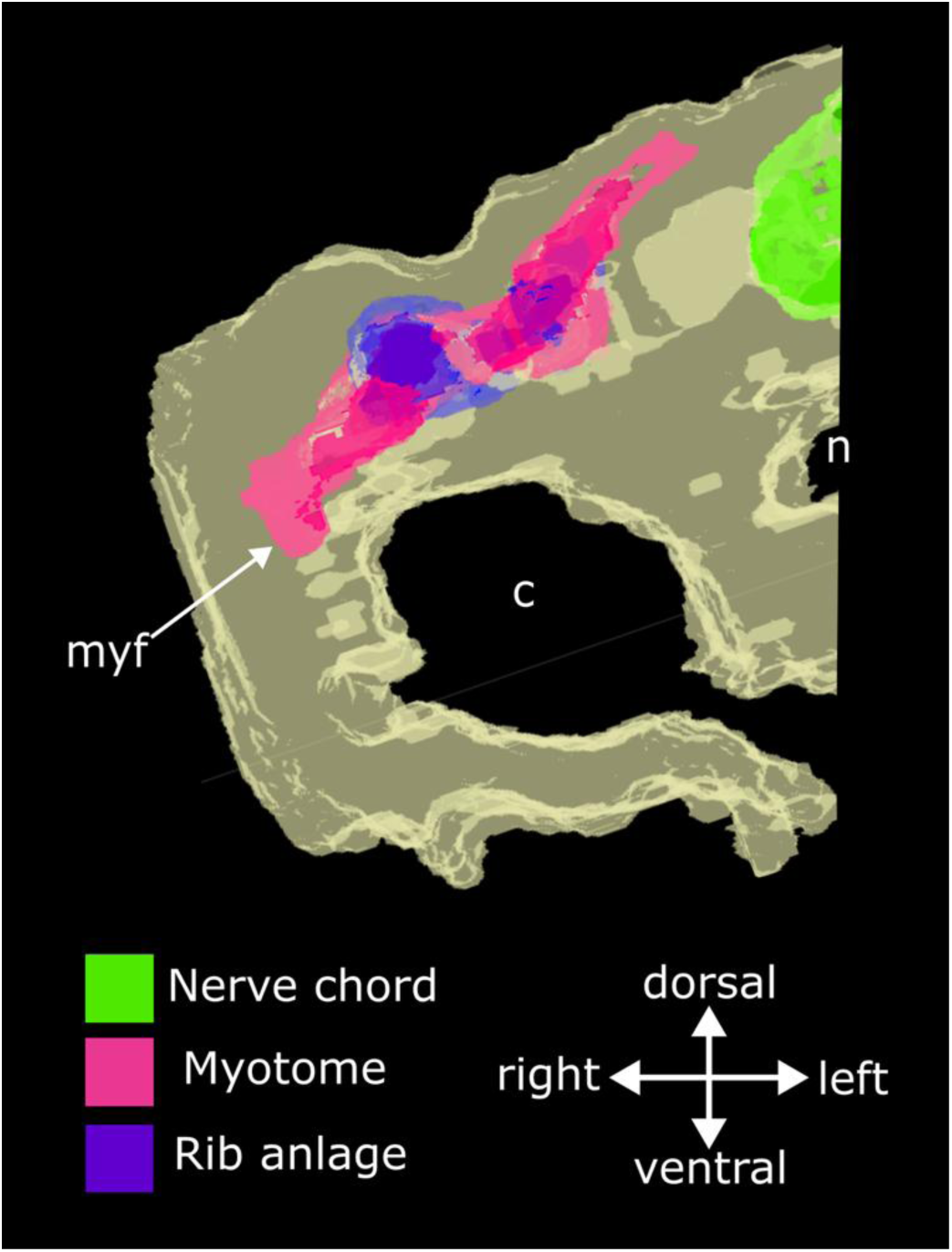
Reconstruction of G12 embryo in coronal cross-section of trunk. G12 cross-section reveals that myotome resides dorsal to the coelom. **Abbreviations:** c, coelom; myf, myotomal front; n, notochord.

**Supplementary Fig 5.**
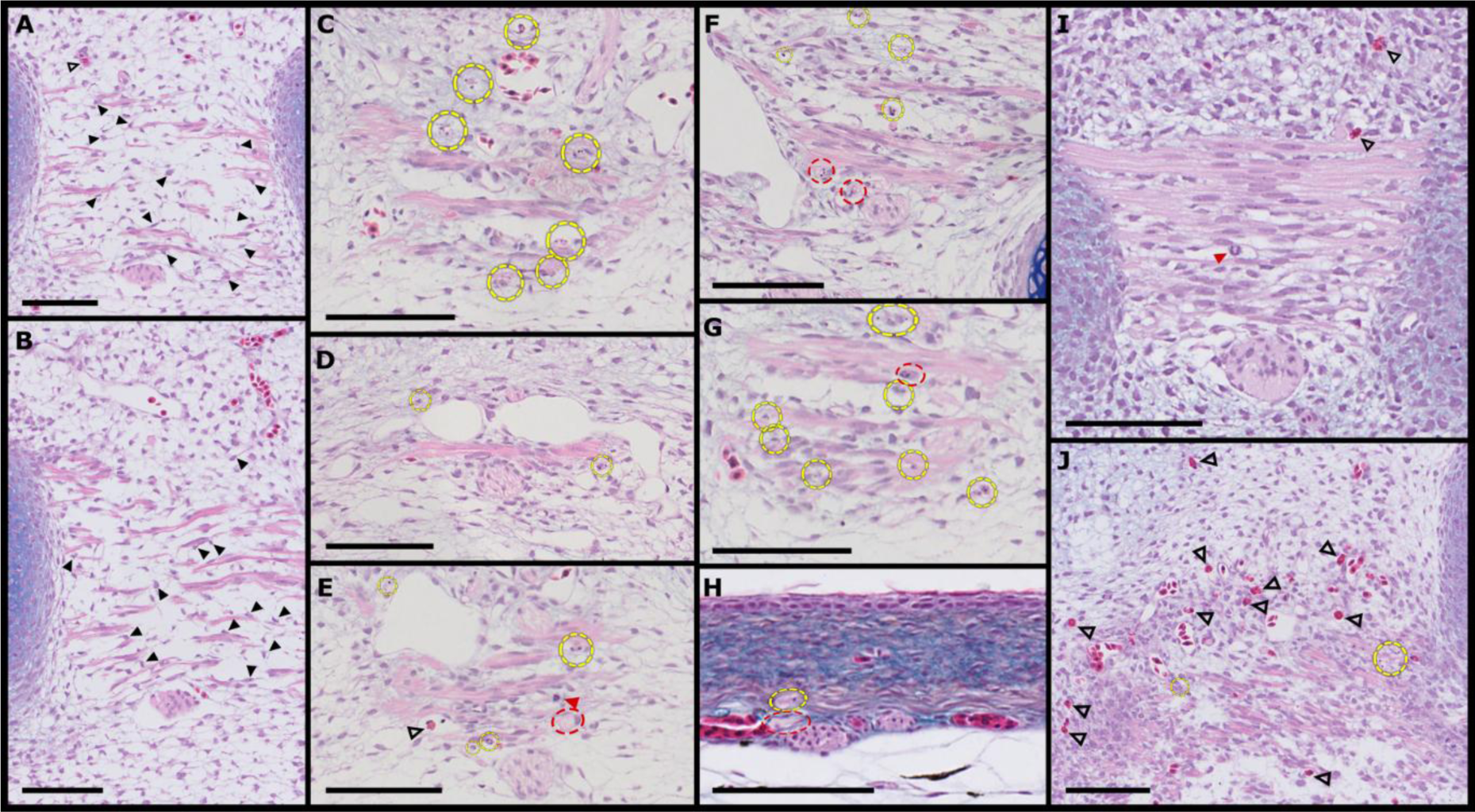
H&E and Alcian blue histology demonstrating mesenchymal infiltration, apoptotic remodeling, and innate immune-associated tissue clearance in turtle intercostal muscle (A,B) G16 sections showing infiltration of stellate mesenchymal cells extending thin cytoplasmic projections toward disorganizing ICM fibres. Mesenchymal cells localize along the periphery of detached muscle fibers. (C,D) G19 sections demonstrating abundant apoptotic bodies within the ICM accompanied by increased vascularization. (E–H) Progressive phagocytic clearance of apoptotic debris by macrophage cells in G19 (E–G) and G20 (H). (I,J) Accumulation of eosinophils within the degenerating ICM, consistent with a localized innate immune and pro-remodeling response associated with cytokine release and angiogenic activation. Black arrowheads indicate invading mesenchymal cells; outlined arrowheads indicate eosinophils; red arrowheads indicate heterophils; yellow dashed circles indicate apoptotic bodies; red dashed circles indicate macrophages. Scale bars represent 100 µm.

#### Trunk muscle volume

Trunk muscle volume was defined as the summed volume of all voxels classified as muscle within the ROI. Voxel-wise tissue classifications were converted to physical volumes in mm³ using the native voxel dimensions:

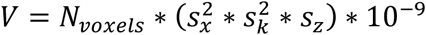

where s_x_ corresponds to the in-plane resolution, s_k_ to downscaling factor applied to x and y, and s_z_ corresponds to the section thickness of 4 µm.

#### Relative muscle volume

Relative muscle volume was defined as the volume of trunk muscles within the total body volume. Total body volume was defined as the sum of all tissues within the decapitated body ROI and all volumes were computed in mm³ using voxel-based conversion as described above.

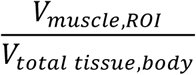

This normalization controls for differences in embryo size between species and developmental stages, enabling comparison of relative muscle growth or loss across developmental time.

To distinguish regional trunk musculature morphogenesis from systemic muscle growth, muscle outside the ROI (e.g., limb and tail musculature) was calculated as the difference between total body muscle volume and ROI muscle volume and normalized to total body volume. This enabled direct comparison between trunk musculature and non-trunk musculature as a function of organism growth across both species.

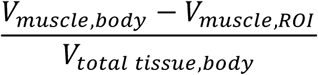

#### β values

To quantify how muscle growth scales with overall embryo size, we modeled the relationship between muscle volume and total body volume using a power-law framework. Allometric scaling relationships were modeled as:

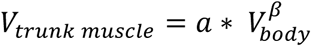

where β is the scaling exponent describing the relative growth rate of muscle with respect to body volume, and *a* is a proportionality constant.

To estimate β, both variables were log-transformed, yielding a linear relationship:

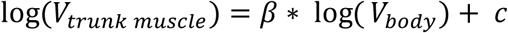

where c = log(a).

Linear and cubic regressions were then performed in log–log space using least-squares fitting and β was calculated as the slope of the fitted linear lines. These equations and transformations allow for estimations and comparisons of tissue scaling exponents across large differences in size and have been utilized for modeling biological systems (Huxley, 1950; Calder, 1984; West et al., 1997)

For mouse samples, a single linear model was used to estimate a global scaling exponent (β_mouse_), reflecting continuous growth dynamics and a R^2^ value was calculated to evaluate fit. For turtle embryos, deviations from linearity in log–log space were observed. To capture these dynamics, two complementary models were applied:

#### Cubic regression

To model the full trajectory of turtle muscle growth and decline, a third-order polynomial was fit to the log-transformed data and a R^2^ value was calculated to evaluate fit:

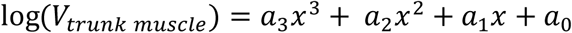

#### Piecewise linear regression (biphasic scaling)

To quantify muscular growth and loss, turtle data were subdivided into early (G12 - G16) and late (G16-G20) intervals. Separate linear regressions were performed on each segment in log–log space:

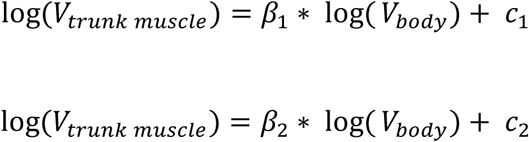

where β₁ and β₂ represent the scaling exponents for early growth and later decay phases, respectively. The overlapping point (TS70/G16) was included in both fits to preserve continuity at the transition.

#### Muscle-skeleton contact

Muscle–skeleton contact (C) was quantified as the number of muscle voxels within a defined volume shell surrounding skeletal tissue. This was calculated by dilating the skeletal mask by 2 voxels followed by intersection with muscle.

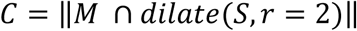

Where M is the binary muscle tissue mask, S is the is the binary skeletal tissue mask and r is the amount the skeletal tissue mask was dilated to calculate intersection with muscle.

This value of muscle contact was normalized (C_norm_) by total skeletal surface area to yield a size-independent muscle-skeleton interface metric allowing for comparisons across different sized embryos.

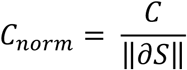

Where ∂*S* is the skeletal boundary voxels identified using 26-neighborhood connectivity.

#### Muscle Packing Fraction and Porosity

An ideal muscle mask representing the spatial extent of the trunk musculature with no holes or infiltrating tissue was generated using closing of the muscle region. This occurred by dilation the raw muscle mask by 1 pixel followed by erosion of the mask by 1 pixel of any non-filled/contacted muscle tissue to fill small intra-muscular gaps and smooth the boundaries of fragmented fibers. Voxels corresponding to skeletal elements and neural tissue were excluded producing a bounding region that captures the expected anatomical domain of the muscle domain. Muscle packing fraction was then calculated as the volume of muscle within the ROI mask intersecting with the idealized mask divided by the idealized mask volume. This measure is analogous to tissue density and provides a spatial proxy for muscle compaction and atrophy.

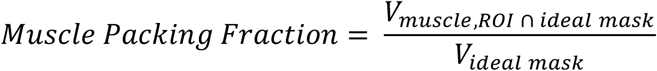

Porosity was then calculated as the fraction of the muscle domain not occupied by muscle tissue (i.e., occupied by extracellular matrix or empty space) defined as:

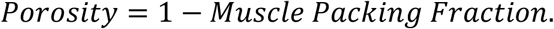

#### Muscle fragmentation

Muscle fragmentation was quantified as the number of spatially disconnected muscle components reflecting the continuity of the muscle tissue. This approach models developing musculature arising from a single, contiguous myotome front and later developing into individual independent epaxial and hypaxial groups (Buckingham, 2001; Cinnamon et al., 1999; Hollway & Currie 2005). Discrete muscle fragments reflect disruption of the normal continuity of the myofiber network and fascicle organization. Increased fragmentation is consistent with muscle atrophy, in which thinner fibers and loss of organized contractile architecture accompany tissue disorganization and fiber loss (Mastaglia & Walton, 1971; Schiaffino et al., 2013).

#### Muscle Surface Area

Muscle surface area was calculated using a triangulated isosurface mesh generated from muscle in the ROI domain downsampled to 15% of its original resolution using nearest-neighbor interpolation. The resulting mesh consists of vertices and triangular faces representing the outer surface of the muscle domain, thus reducing sensitivity to voxel-level irregularities. Vertex coordinates were rescaled to physical units (mm) using the original voxel spacing in the x, y, and z dimensions to account for downsampling. Surface area was then computed as the sum of the areas of all triangular faces in the mesh, calculated using the cross product of edge vectors for each triangle:

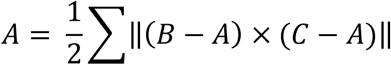

where A, B, C are the vertices of each triangular face.

#### Developmental Time Normalization

Developmental time was normalized relative to total gestation, with a conserved developmental time point in each species serving as the reference point (t = 0) for comparative analyses. For this study, the conserved/phylotypic stage was defined as E9.5 in mouse (Irie & Kuratani, 2011) and Greenbaum stage G11 in turtle (Wang et al., 2016), reflecting genomic and morphological conservation across species along with our own assessment of conserved limb and trunk characters. To align with experimental measurements, the phylotypic stage was converted to days post-fertilization, yielding t_phylo_= 9.5 days for mouse and t_phylo_= 8 days for turtle (Noravian et al., 2025). Subsequent developmental time points (t_i_) were normalized as the fraction of gestation remaining after the phylotypic stage (t_norm_):

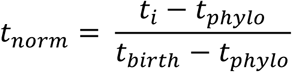

Where t_birth_ represents species-specific gestation length (mouse: 20 days (Murray et al., 2010); turtle: 60 days (Noravian et al., 2025)). This normalization accounts for interspecific differences in both early and late gestation, enabling comparisons of morphogenesis between mouse and turtle embryos. All normalized developmental times were reported on the x-axis of line plots.

